# Differential reliance on sensory reinstatement and internally transformed representation during vivid retrieval of visual and auditory episodes

**DOI:** 10.1101/2025.06.16.659408

**Authors:** Lei Zhang, Claude Alain, Bradley R. Buchsbaum

## Abstract

Auditory memory is considered less detailed yet more durable than visual memory, implying a modality-specific memory retrieval process. We used fMRI and multivoxel pattern analyses to examine how 25 participants encoded and retrieved naturalistic sounds and videos. Both auditory and visual targets reinstated item-specific fine activation patterns in the association cortex during retrieval, and reinstatement strength correlates with subjective memory vividness. However, auditory episodes showed a markedly larger reliance on internally constructed representations than visual episodes, quantified by retrieval-retrieval similarity after removing encoding traces. Sensory reinstatement correlated more to the (detail-related) posterior hippocampus, while internal representations also correlated to the (gist-related) anterior hippocampus. Furthermore, temporal voice areas preserved gist-level (human versus non-human) information from encoding to retrieval, whereas fusiform face representations degraded. These findings reveal that auditory and visual memories share a common sensory reinstatement mechanism, but differ in the neural mechanism that supports retrieval, with participants favouring gist over perceptual details during auditory memory retrieval.

## Introduction

Vision and audition are the chief gateways through which we learn about—and adapt to—the world. Behavioural studies show a clear asymmetry between the two: long-term memories for sounds tend to discard fine perceptual detail and preserve the overarching “gist,” whereas visual memories more often retain far richer specifics^1–3^. Therefore, clarifying how the brain retrieves auditory versus visual events is essential for a general theory of long-term memory. Contemporary models posit that successful recall hinges on neural pattern reinstatement—the re-emergence, voxel by voxel, of the activity pattern originally present at encoding^4–9^. While this theory has been well-established in visual episodic memory research, it remains to be seen whether episodic memory retrieval from other modalities, such as auditory episodic memory, engages the same neural reinstatement mechanisms or employs distinct strategies.

Evidence from neuroimaging studies suggests that auditory and visual processing are hierarchically organised in the cortex^10,11^. Primary areas in both systems process basic perceptual features, such as frequency and loudness for auditory stimuli and line orientation and brightness for visual stimuli. The higher-order auditory and visual cortices encode abstract and complex information, such as object identity^10,11^. Although recalling a visual scene reliably reactivates the higher-order visual cortex, it is still uncertain whether the same reinstatement mechanism underlies memory retrieval in other domains, such as audition. Evidence from functional magnetic resonance imaging (fMRI) studies suggests that recalling auditory events reactivates association areas involved during encoding^12–14^. However, it is still unknown whether the fine-grained spatial patterns evoked when we recall auditory events resemble those present during their encoding, as has already been demonstrated for visual memory. Moreover, the similarities and differences between cortical reinstatement of auditory and visual scenes have yet to be identified. Because auditory and visual perception have overlapping circuitry and are hierarchically organised, we predict that recalling sounds would involve neural pattern reinstatement comparable to that observed during visual retrieval. Yet behavioural evidence shows that auditory memories, while longer-lasting, are coarser and more gist-driven than their visual counterparts—hinting that auditory retrieval may depend on mechanisms that diverge from those supporting visual recall^1–3^.

In this two-session fMRI study, we used an episodic “pattern completion” paradigm to investigate auditory and visual episodic memory retrieval cued by audio or video clips. Participants completed both encoding and retrieval runs. During encoding, participants studied sequential pairs of stimuli—everyday sounds and/or videos—in which the first item served as the cue and the second as the target to be remembered. Retrieval runs used a cue-recall task to test auditory or visual memory (Fig. 1A, B). In Session 1, the pairs were entirely new to participants; by Session 2, all cue-target associations had been fully learned.

**Figure 1.**
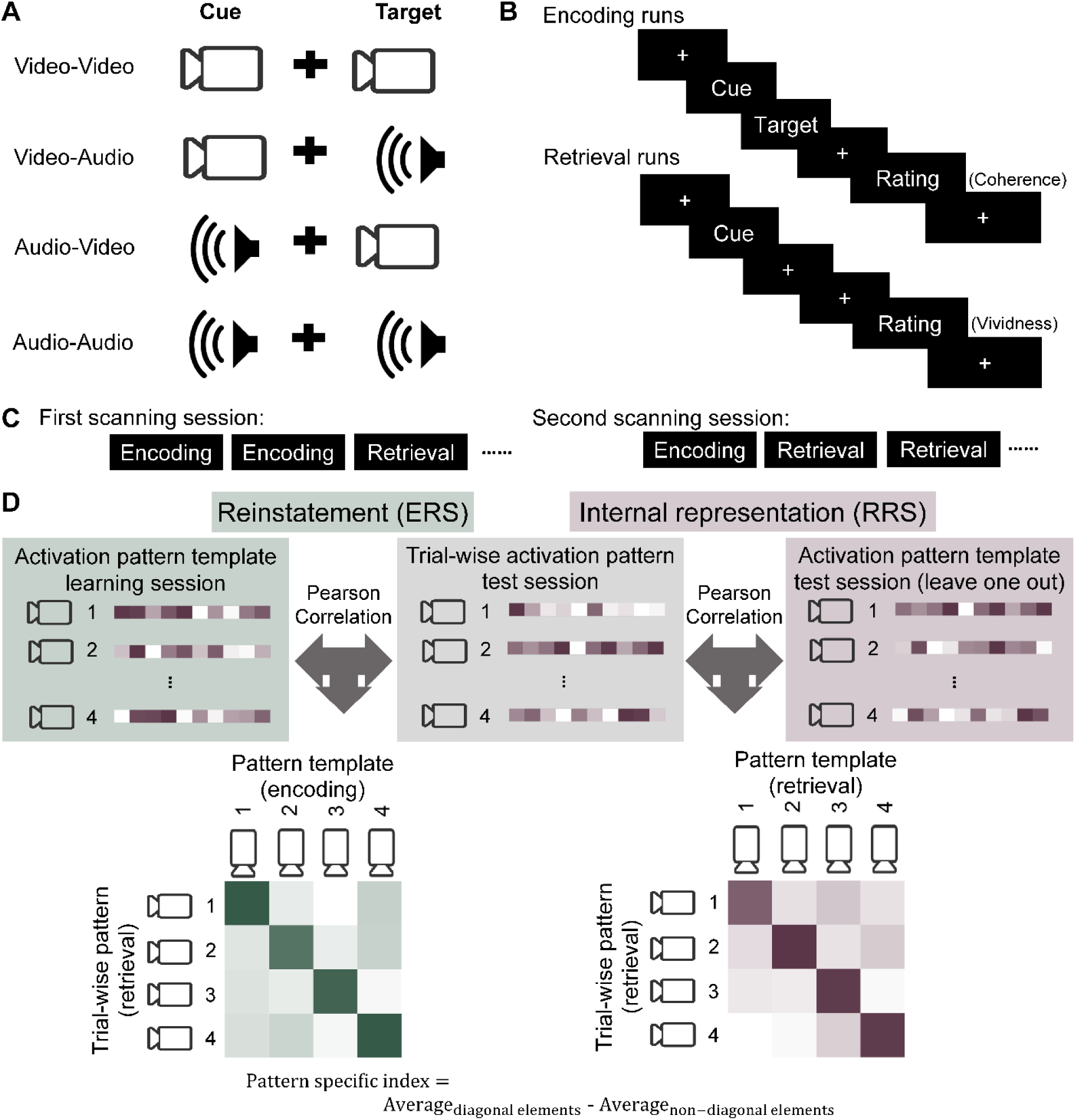
Paradigm and analysis pipeline. (A) Two sequential clips formed a paired associate. (B) In encoding blocks, paired associates were presented once, and participants rated for cue-target coherence. Retrieval blocks began with presentation of the cue, followed by silent recall of the target, and participants gave a vividness rating. (C) Each scanning session comprised alternating encoding and retrieval blocks. (D) ERS and RRS PSI calculation. Trial-wise retrieval data were correlated to the template pattern of items during encoding (ERS) or retrieval (RRS). PSI was defined as the within-item correlation minus the mean between-item correlation. ERS, encoding-retrieval similarity; RRS, retrieval-retrieval similarity; PSI, pattern specificity index.

We used multivoxel pattern analysis (MVPA) to determine which items or stimulus features were encoded in the observed brain-activity patterns^15^. Encoding–retrieval similarity (ERS) quantifies how closely voxel-wise activation patterns during retrieval match those recorded at encoding, providing a direct measure of neural reinstatement of the original memory trace^16^. Evidence indicates, however, that retrieved traces may be transformed and are not exact replays of the encoding pattern^17–19^. Retrieval-retrieval similarity (RRS) tracks how consistently internally generated representations recur across retrieval trials, regardless of whether they faithfully match the original encoding pattern or have been transformed. ERS and RRS draw on different correlation matrices, encoding-retrieval for ERS versus retrieval-retrieval for RRS, so they are statistically independent. However, they can coincide in regions that reinstate sensory traces and maintain transformed memories^20^. We isolate item-specific patterns by computing a pattern specificity index (PSI)—the difference between same-item and different-item ERS or RRS values^15^. Memory reinstatement and representational content can also be explored with multivoxel pattern classification analysis^16^. To enhance the robustness of our findings, we report both measures, which are expected to align. Given that auditory memory tends to be less detailed and gist-based than visual memory^1,2^, we hypothesize that auditory memory would exhibit a stronger reliance on an internally constructed or transformed representation when compared to visual memory.

Critically, the hippocampus is recognized as a central hub for episodic memory in the visual domain^21,22^, and it has also been implicated in processing simple auditory sequences^23^. Yet the precise role of hippocampal subregions in retrieving detailed vs. gist-like auditory episodes remains underexplored. A proposed gradient along the hippocampal long axis suggests that the posterior hippocampus preferentially supports the retrieval of rich perceptual details. In comparison, the anterior hippocampus is more strongly associated with broader, gist-like representations^22^. Along the hippocampal long axis, we predict that the posterior hippocampus will correlate more strongly with sensory-driven ERS. However, the anterior hippocampus should correlate more strongly with transformed or reconstructed representations (RRS).

We found that auditory and visual memories are both reinstated in their respective sensory association cortices during retrieval, and activity in these regions scales with trial-by-trial ratings of vividness. Yet auditory recall leans more heavily than visual recall on internally constructed, gist-like representations. Sensory-based reinstatement couples exclusively with the posterior hippocampus, which is tuned to perceptual detail, whereas these internally generated representations engage both anterior and posterior segments, implicating the hippocampus in supporting both gist and detail. Lastly, anterior hippocampal activity correlated to auditory and visual memory vividness ratings. These findings suggest partially distinct neural mechanisms support auditory and visual episodic memory retrieval.

## Results

### Auditory episodic memory is less vivid than visual episodic memory

Vividness ratings increased with stimulus repetitions in the first scanning session (F_run_ (2,46) = 112.46, p < 0.001). In the second scanning session, although the effect of run number was significant: F_run_ (5,22) = 3.24, p = 0.024, post-hoc pairwise tests showed no specific run-to-run differences (|t(22)| < 2.853, |p_fdr_| > 0.077). Thus, vividness was already high and stable throughout session 2—most likely because the additional pre-session training had brought all cue-target pairs to ceiling memory strength.

Overall, vividness ratings were higher for video than audio targets, regardless of the cue modality (First session: F_target_ (1, 23) = 56.11, p < 0.001; Second session: F_target_ (1, 22) = 31.236, p < 0.001). These findings suggest that visual episodic memory is more vivid than auditory memory, consistent with previous behavioural results^1–3^.

### Parsing retrieval-specific neural activity through differential target modality subtraction

One of the main challenges in a pattern completion paradigm is separating cue-related BOLD responses from true retrieval-related activity. Using auditory retrieval as a baseline for visual retrieval (and vice versa) helps highlight retrieval-specific activity. For instance, activity in visual areas decreases irrespective of the target’s modality after a visual cue, potentially obscuring activation evoked by visual retrieval unless it is compared with an auditory target baseline. In this study, we subtracted the neural activity elicited by the visual target from the activity elicited by the auditory target. Subtracting the visual target from the auditory target, and vice versa controls for the influence of cue modality, which was consistent across both conditions. Specifically, the contrast (Video_cue_ Audio_target_ – Video_cue_ Video_target_) removes the visual cue effect, while the contrast (Audio_cue_ Audio_target_ – Audio_cue_ Video_target_) removes the audio cue effect. Averaging these differences yields the main effect of the target modality. For example, auditory-item representation or reinstatement measures in auditory areas are calculated as follows:

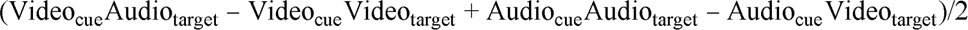

Areas showing significantly positive values indicate that auditory targets show enhanced activity compared to visual targets, after correcting for modality-specific carry-over effects. A similar process is also performed for visual-item measures. We report the corrected measures with the cue effect removed in the following univariate and multivoxel analyses.

### Sensory association areas show increased activities during both auditory and visual memory retrieval

We examined sensory reinstatement during episodic memory retrieval on auditory and visual cortex regions of interest (ROIs) (Figure 2b) using retrieval data from the second session, as participants had fully memorized the pairs by then. We observed higher activation in auditory association ROIs during auditory memory retrieval and increased activation in visual association ROIs during visual memory retrieval (Figure 2C, D, p_fdr_ < 0.05). In comparison, primary auditory and visual areas did not display memory-related activation for either cue modality (Figure 2C). In primary visual areas, activation decreased during visual memory retrieval (Figure 2C, D). The small memory-related activity in early sensory areas aligned with previous studies as no external sensory stimuli were presented during the retrieval phase^13,24–26^. Higher-level sensory areas, where more abstract features are processed, showed activation during mnemonic retrieval, highlighting the distinction between internal memory signals and external stimuli.

**Figure 2.**
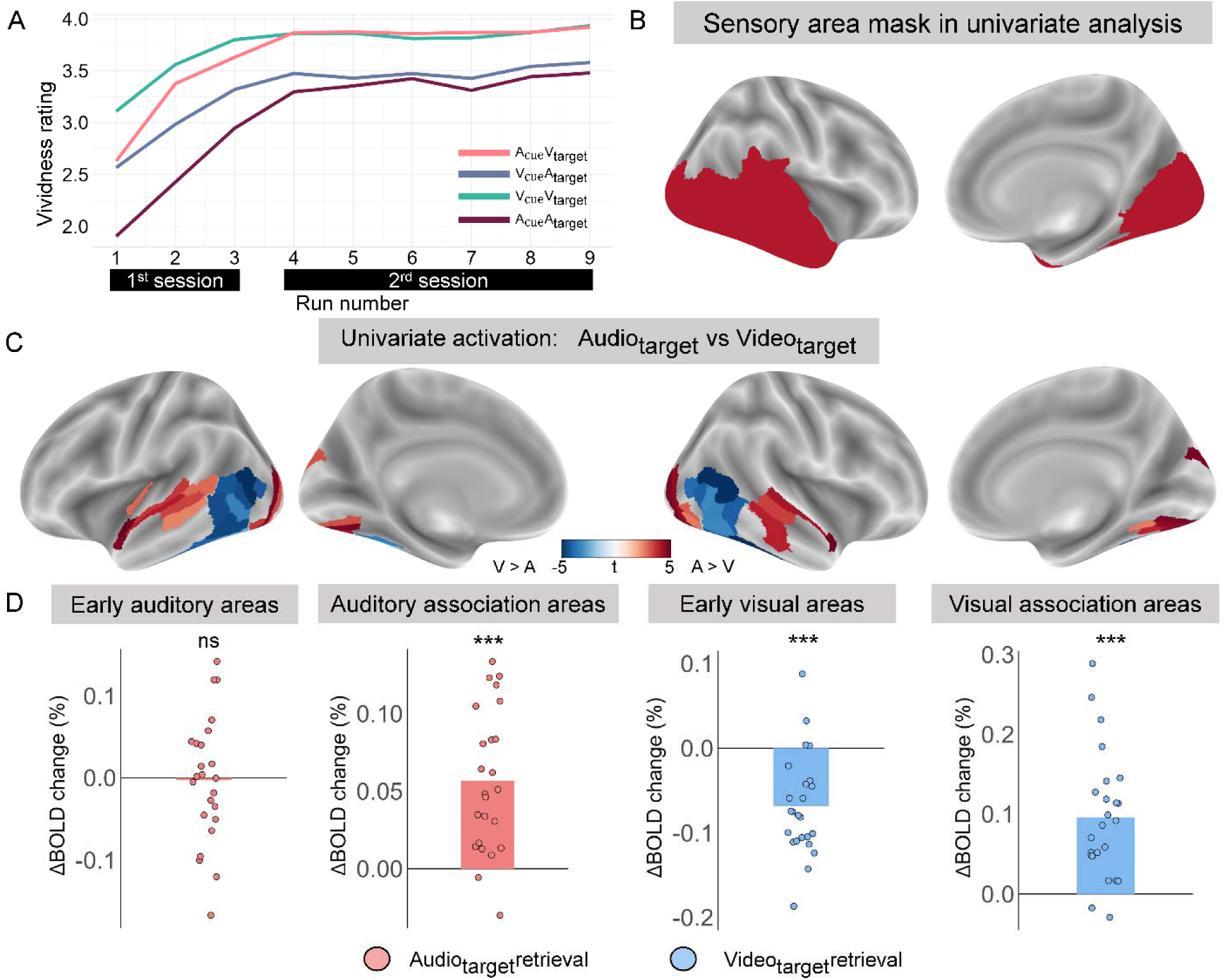
Behaviour and univariate analysis results. (A) The averaged vividness rating of four conditions across nine retrieval runs. (B) Anatomical sensory areas mask used in univariate analysis. (C) ROIs showed significant difference of BOLD activation between auditory and visual memory retrieval (p_fdr_ < 0.05). (D) During auditory retrieval, auditory association areas show increased activity while early auditory areas do not. Visual association areas showed increased activities during visual retrieval while early visual areas showed decreased activity. ***P < 0.001, ns, not significant.

### Auditory and visual items are both reinstated in corresponding association areas

We examine sensory reinstatement during memory retrieval using MVPA. Based on the results of the univariate analysis, we anticipated that only sensory association areas would show significant reinstatement. We identified four clusters (Figure 3A, table S3, early auditory cortex, auditory association areas, early visual cortex, and visual association areas) from the results of the univariate analysis. Left and right ROIs were averaged since lateralization is not an interest in this study (for details, see methods). We conducted two complementary analyses: (1) we computed ERS to test whether the retrieval activation pattern resembled the encoding pattern as indexed by the PSI (Figure 1D, for details, see methods), and (2) Multivoxel pattern classification analysis, which involved training a classifier on encoding trials to identify specific retrieved target stimuli at retrieval. Both analyses test whether items are represented in sensory regions during retrieval in a manner resembling their representations at encoding, i.e., sensory reinstatement. High classification accuracy indicates items are well-differentiated in a given region, while high PSI shows stable and distinct spatial patterns evoked by different stimuli.

**Figure 3.**
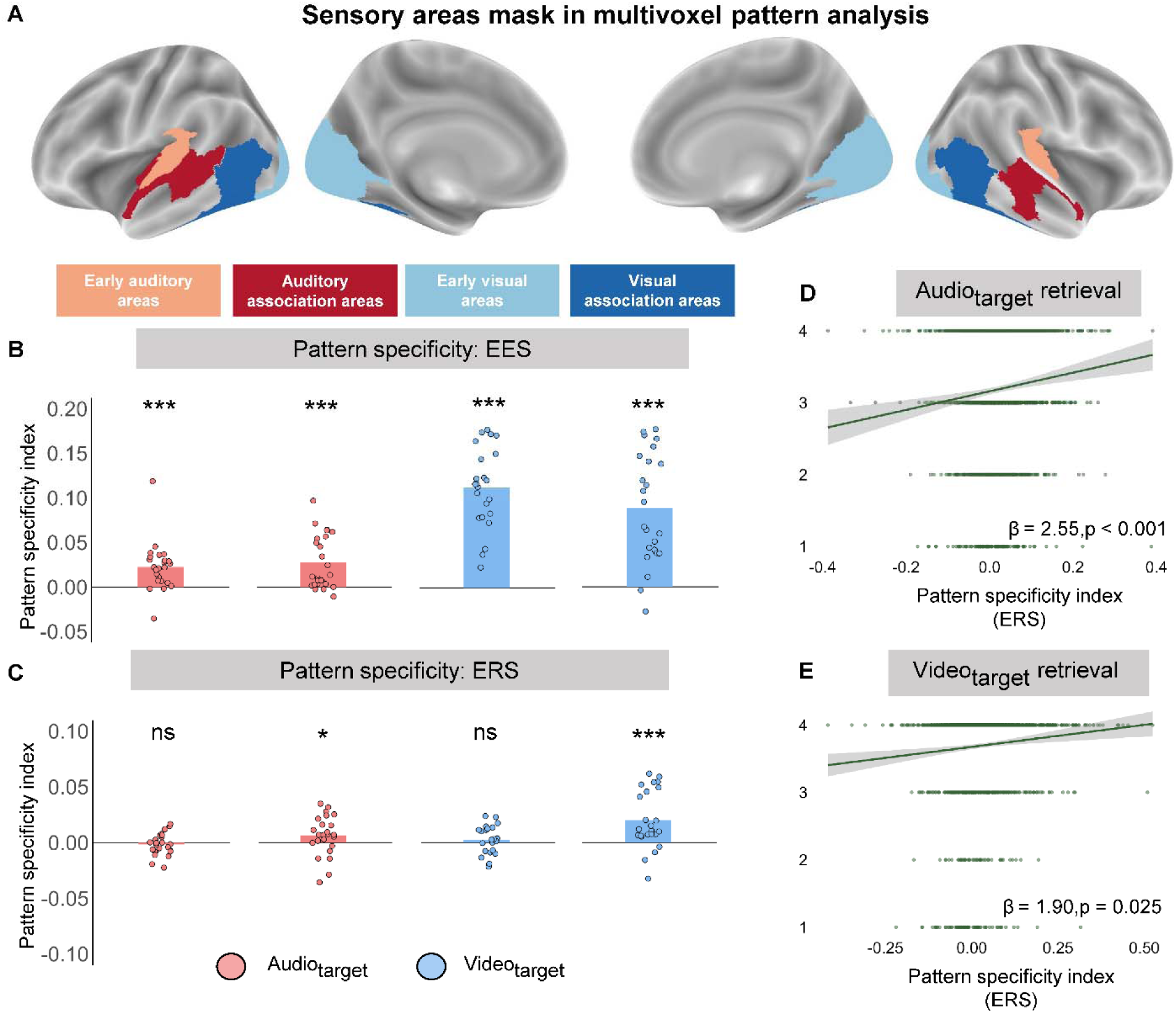
Neural reinstatement in sensory association cortex. (A) Four clusters in sensory areas used for MVPA: early auditory areas, auditory association areas, early visual areas, and visual association areas. (B) All four clusters showed robust representation of stimulus items during encoding. (C) Only visual and auditory association areas showed robust reinstatement of stimulus items during retrieval. (D, E) ERS correlated with vividness ratings for their respective modalities in both auditory and visual association areas. *P < 0.05, ***P < 0.001, ns, not significant. ERS, encoding-retrieval similarity; EES, encoding-encoding similarity.

First, we performed encoding-encoding similarity (EES) and classification analysis during encoding runs from both sessions to show that the items are represented in sensory clusters (for details, see methods). Both early and auditory association areas showed a significant representation of auditory items, and both early and visual association areas showed a significant representation of visual items (Figure 3B, S1A, and Table S1). Next, we tested for sensory reinstatement in sensory ROIs. Both complementary measures yielded consistent results. Significant sensory reinstatement was found in sensory association areas (Figure 3C, S1B, and Table S1). No evidence of reinstatement was found in early sensory areas (Figure 3C, E, and Table S1). To test whether more robust sensory reinstatement correlates with increased memory vividness during learning, we used a cumulative link mixed model^27^ to assess the trial-wise relationship between sensory reinstatement and memory vividness. We included first-session data to capture a broader range of vividness ratings. Both PSI and classification probability (for the “true” item) positively predicted vividness ratings when controlling for stimulus type and cue modality (Figure 3D, E, S1C, D). This suggests that sensory reinstatement measure could be a potential objective index for memory vividness regardless of sensory modality. From all these results, we found that auditory memory retrieval shares a common sensory reinstatement process with visual memory retrieval, which correlated with memory vividness.

### RRS: Auditory memory retrieval showed greater reliance on internally constructed representations than visual memory retrieval

Next, given that retrieval does not simply mirror encoding activity^18,28^, we tested whether memory items are robustly represented as their own form instead of just reinstatement of memory encoding. Also, we asked whether visual and auditory memory retrieval showed different biases for internal representation and reinstatement of item encoding. We used a leave-one-run-out cross-validation procedure to examine the internal memory representation to conduct classification and RRS analysis, assessing the consistency of spatial activity patterns during memory retrieval of the same item (for details, see methods).

We found consistent results in memory representation. Auditory and visual memory are represented in sensory association areas (Figure 4A, S2A, and Table S1). No evidence of memory representation was found in early sensory areas (Figure 4A, S2A, and Table S1). These results suggest that sensory association areas maintain stable internal representations of items during memory retrieval, and early sensory areas do not reliably reinstate robust representations of the recalled item. PSI also positively predicted vividness ratings when controlling for stimulus type and cue modality (Figure 4B), while classification probability only predicted auditory vividness (Figure S2B).

**Figure 4.**
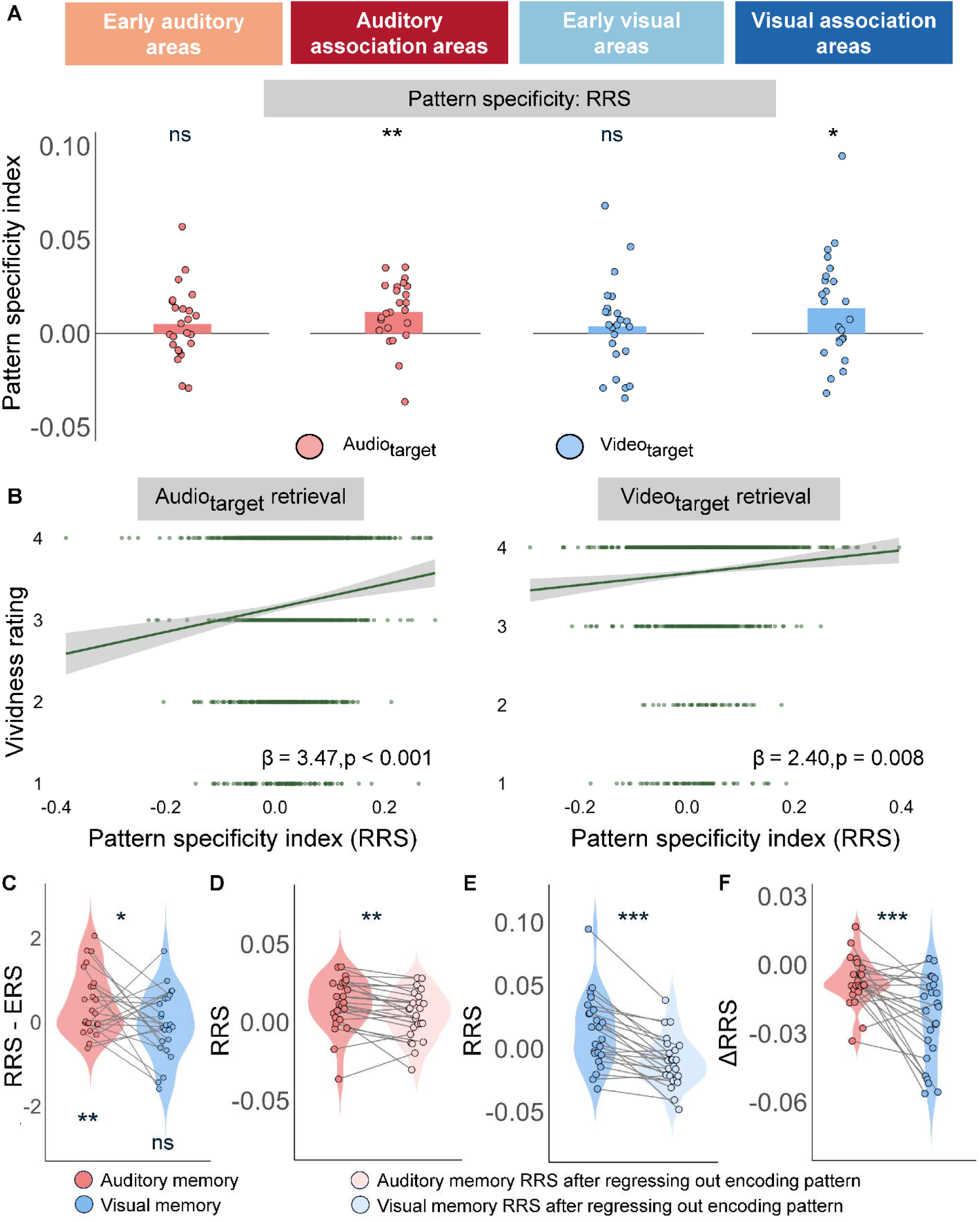
Internal representation in sensory association cortex. (A) Only visual and auditory association areas showed a robust internal representation of stimulus items during retrieval. (B) Auditory and visual RRS correlate with vividness rating in auditory and visual association areas. (C) Auditory memory showed a higher bias for internal representation than visual memory. (D, E) Regressing out trial-wise encoding patterns (to isolate activity unique to retrieval) reduced RRS for both modalities. (F) This reduction was smaller for auditory items in auditory association areas than for visual items in visual association areas. *P < 0.05, **P < 0.01, ***P < 0.001, ns, not significant. RRS in visual association areas. ERS, encoding-retrieval similarity; RRS, retrieval-retrieval similarity.

However, our leave-one-run-out classification and RRS metrics might still be contaminated by reinstatement: if an item shows high encoding–retrieval similarity, that same overlap could artificially inflate the correlations we compute solely among retrieval patterns. Therefore, we defined internal-representation bias as the difference between representation and reinstatement. A greater bias for internal representation indicates that memory retrieval relies more on internal representation. We found that auditory memory showed a greater bias for internal representation while no bias for internal representation was observed for visual memory (Figure 4C, Auditory memory: t_PSI_ (23) = 2.99, p_PSI_ = 0.007; Visual memory: t_PSI_ (23) = -0.33, p_PSI_ = 0.741). Similar results were found with the classification method. Auditory memory showed a marginally significant bias for internal representation, while visual memory showed a marginally significant bias for reinstatement (Figure S2C, Auditory memory: t_accuracy_ (23) = 1.74, p_accuracy_ = 0.095; Visual memory: t_accuracy_ (23) = -1.74, p_accuracy_ = 0.095). To investigate whether auditory memory relies more on internal representation to reinstatement than visual memory retrieval, we compared the bias to internal representation between the two modalities. We found that auditory memory showed a higher bias for internal representation than visual memory (Figure 4C, S2C, t_PSI_ (23) = 2.16, p_PSI_ = 0.041, t_accuracy_ (23) = 2.36, p_accuracy_ = 0.027).

Next, in addition to subtracting the reinstatement measure from the internal-representation measure to control for the influence of reinstatement, we performed a “reinstatement-free” RRS analysis. We regressed the encoding-phase template pattern of each item out of every retrieval trial to obtain residual activation patterns that did not contain encoding information. We then applied the same RRS analysis to these residuals. As expected, the RRS measure decreased after removing the reinstatement component for visual targets, using both PSI and classification accuracy measures (Figure 5E and Figure S2E, t_PSI_ (23) = -6.30, p_PSI_ < 0.001, t_accuracy_ (23) = 6.77, p_accuracy_ < 0.001). For auditory targets, the RRS measure decreased after removing the reinstatement component when assessed with PSI, but not with the classification accuracy measure (Figure 5D and Figure S2D, t_PSI_ (23) = -3.39, p_PSI_ = 0.002, t_accuracy_ (23) = 1.24, p_accuracy_ = 0.226). The adverse effect of removing the reinstatement component was significantly smaller for auditory-item RRS in auditory association areas than for visual-item RRS in visual association areas (Figure 4F and Figure S2F, t_PSI_ (23) = 3.95, p_PSI_ < 0.001, t_accuracy_ (23) = 4.59, p_accuracy_ < 0.001).

**Figure 5.**
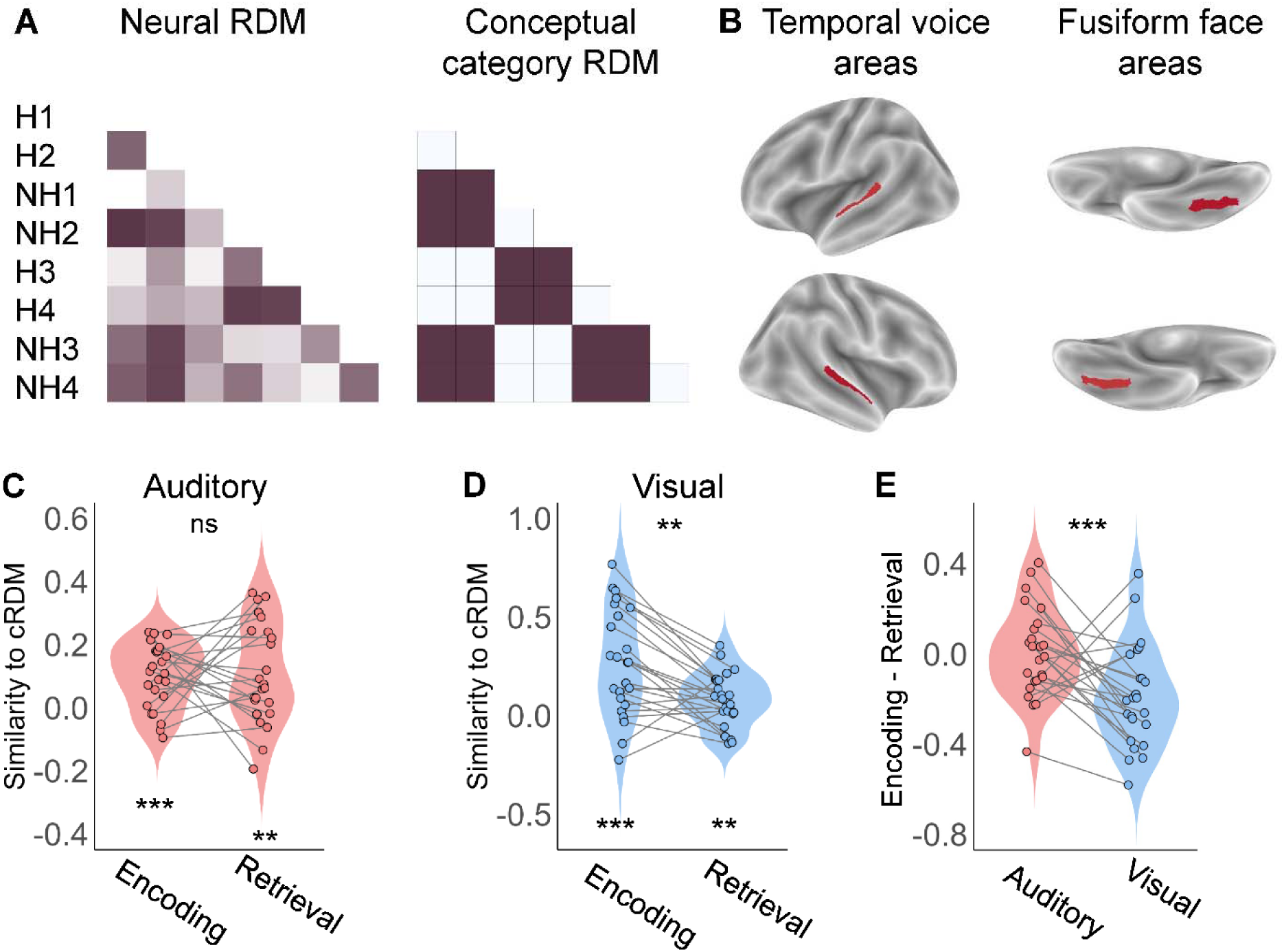
Category representation in the temporal voice area and the fusiform face areas. (A) Neural RDM and conceptual category RDM. (B) Left and right temporal voice area and fusiform face areas. (C) Auditory category representation is robust in the temporal voice area during encoding and retrieval, with no difference between phases. (D) Visual category representation is robust in fusiform face areas during encoding and retrieval, but it decreases in the retrieval phase compared with encoding. (E) The drop in auditory category representation from encoding to retrieval in the temporal voice area is significantly smaller than the corresponding drop in visual category representation in the fusiform face areas. **P < 0.01, ***P < 0.001, ns, not significant.

These results reveal similar sensory reinstatement and representation in the sensory association cortex during auditory and visual memory retrieval. They also showed differences between the two sensory modalities. Auditory memory retrieval depends more on internally constructed representations than visual memory retrieval. The latter is consistent with behavioural studies of auditory memory, which suggest that it is more gist-based^1^. That is, the pattern of brain activation provides converging evidence that self-constructed internal representation is the dominant mode for auditory memory retrieval.

### Categorical RSA: auditory memory preserves gist-level representation

RRS results indicated that auditory memory retrieval showed an advantage in internally constructed representation, suggesting that auditory memory is more gist-based. We, therefore, carried out a categorical RSA to assess gist-based representations (human versus non-human) during encoding and retrieval in the temporal voice area and fusiform face area (Fig. 5A, B).

We found that both the temporal voice and fusiform face areas showed robust human versus non-human representations during encoding and retrieval (Figure 5C and D, Table S2). In the fusiform face area, category representation of visual items during retrieval significantly decreased compared to that during encoding (Figure 5D, Table S2). By contrast, in the temporal voice area, the category representation for auditory items during retrieval was as strong as during encoding (Figure 5C, Table S2). When we contrasted the drop in representation strength from encoding to retrieval, the decline was markedly smaller for auditory gist in the temporal voice area than for visual gist in the fusiform face area (Fig. 5E). This indicates that gist-level representations of sounds survive the perception-to-memory transition better than those of images, offering a neural account for the greater durability of auditory gist observed behaviourally^2^.

### Hippocampal activity tracks the vividness of auditory and visual memories

The hippocampus is crucial for episodic visual memory^21,22^. Here, we asked about the role of the hippocampus in auditory and visual memory retrieval. We separated the hippocampus into anterior and posterior parts using Melbourne Subcortex Atlas^29^. We correlated voxel-wise hippocampal activity with each participant’ memory vividness of each trial. We observed higher anterior hippocampal activity associated with greater memory vividness for auditory and visual targets (Figure 6A–F), whereas the posterior hippocampus showed no such effect. For auditory targets, the left anterior hippocampal activity correlated positively with memory vividness (left: t(23) = 2.82, p_fdr_ = 0.039; right: t(23) = 1.20, p_fdr_= 0.482) instead of the posterior hippocampus (left: t(23) = -0.38, p_fdr_ = 0.706; right: t(23) = 0.40, p_fdr_ = 0.706). For visual targets, left and right anterior hippocampal activity correlated positively with memory vividness (left: t(23) = 3.27, p_fdr_= 0.013; right: t(23) = 2.96, p_fdr_ = 0.014) instead of the posterior hippocampus (left: t(23) = 2.17, p_fdr_ = 0.054; right: t(23) = 1.34, p_fdr_ = 0.193). A voxel-wise illustration confirmed that the anterior hippocampus showed a higher correlation than the posterior part (Figure 6B, C, E, F). We also performed linear regression analysis to test the relationship between the correlation coefficients of each voxel and the y (posterior-anterior) and z (inferior-superior) axis coordinates to confirm that further. We found the more inferior and anterior voxels showed higher correlation coefficients (all |t| > 5.697, all p_fdr_ < 0.001). However, no significant correlation was found between sensory reinstatement and hippocampus activity.

**Figure 6.**
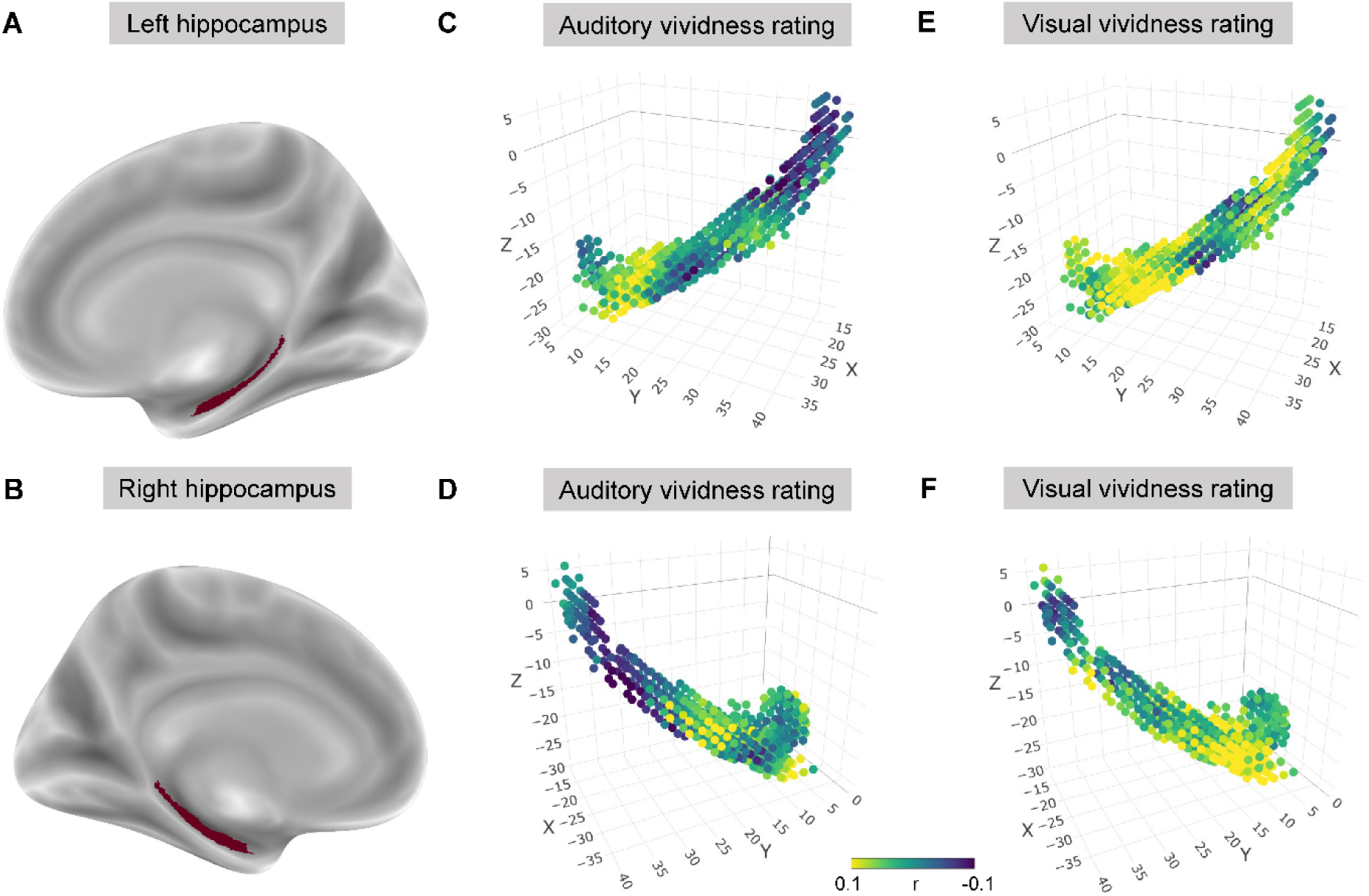
Anterior hippocampal activity predicted vividness rating. (A, B) Anatomical mask of the left and right hippocampus. (C-F) The anterior hippocampus is more correlated to the auditory and visual vividness rating.

### Hippocampal-cortical informational interaction analysis: The anterior hippocampus interacts with internal representation while the posterior hippocampus interacts with reinstatement

Next, we investigated the informational interaction between hippocampus subfields and sensory association cortex by correlating trial-wise reinstatement and representation measures using a linear mixed model (for details, see methods). We speculate that visual memory is more detailed, which explains the stronger bias for reinstatement—reflected in greater neural similarity to encoding activity when perceiving item details. Building on the proposal that gist and detailed representations are organised along the hippocampus’s longitudinal axis^22^, we hypothesized that reinstatement would involve the posterior hippocampus, while representation would engage the anterior hippocampus. Given that hippocampal activity is similarly associated with vividness in auditory and visual memory, and our focus on the relationship between ERS and RRS measures and hippocampal subfields, we combined auditory and visual trials for the hippocampal–cortical informational interaction analysis.

Through trial-wise linear mixed effect modelling, we found that reinstatement of sensory association areas interacted with the posterior hippocampus rather than the anterior hippocampus (Table 1, anterior hippocampus: standardized β = 0.012, p = 0.313; posterior hippocampus: standardized β = 0.072, p < 0.001; For separate auditory and visual results, see Table S3). Applying Cumming’s (2009) procedure for comparing dependent regression coefficients (see *Methods*^30^), we confirmed that the posterior coefficient was significantly larger than the anterior one (*p* < 0.05), underscoring a stronger coupling between sensory reinstatement and posterior-hippocampal activity. Next, we found that the strength of the representation in sensory association areas correlated significantly with both the anterior and posterior hippocampus (Table 1, anterior hippocampus: standardized β = 0.077, p < 0.001; posterior hippocampus: standardized β = 0.044, p = 0.008). The anterior hippocampus’s standardized beta coefficient was larger than the posterior hippocampus’s. But the difference between these two standardized beta coefficients did not pass the Cumming (2009)’s significance test. Therefore, these results partially met our expectations. Reinstatement is more related to detailed representations, while internal representations are more related to gist information.

**Table 1.** Hippocampal-cortical informational interaction analysis results. Standardized beta coefficient of anterior and posterior hippocampus predicting cortical memory reinstatement and representation from linear mixed model.

|  | Memory reinstatement |  | Memory representation |  |
| --- | --- | --- | --- | --- |
| Anterior hippocampus |  |  |  |  |
| reinstatement | $B = 0.016$ | representation | $B = 0.077$ | |
| | $P = 0.313$ | | $P < 0.001$ | |
|  | 95% CI: -0.017, 0.050 |  | 95% CI: 0.043, 0.111 |  |
| Posterior hippocampus |  |  |  |  |
| reinstatement | $B = 0.072$ | representation | $B = 0.044$ | |
| | $P < 0.001$ | | $P = 0.008$ | |
|  | 95% CI: 0.041, 0.091 |  | 95% CI: 0.014, 0.076 |  |

## Discussion

This fMRI study dissected how the brain retrieves auditory and visual episodes. Both modalities showed sensory-cortical reinstatement, and the strength of that reinstatement predicted trial-by-trial vividness. Yet several clear asymmetries emerged. Retrieval-retrieval similarity (RRS) uncovered internally generated patterns independent of the original encoding trace; auditory recall relied on these constructive representations more than visual recall. Categorical RSA confirmed that high-level gist information for sounds was better preserved from perception to memory than the equivalent visual gist. Informational-connectivity analysis further revealed a hippocampal gradient: sensory reinstatement coupled selectively with the posterior hippocampus, which supports fine perceptual detail, whereas internally generated patterns engaged both anterior and posterior segments. Across modalities, vividness ratings were more strongly related to anterior than posterior hippocampal activity, consistent with a long-axis gist-to-detail gradient. These results show that auditory and visual memories share a common reinstatement scaffold but diverge in balance: auditory retrieval leans toward durable, gist-based reconstructions, whereas visual retrieval retains richer sensory details.

Auditory and visual perception share some common neural mechanisms. The early sensory cortex processes basic features, such as pitch and amplitude for sounds and colour and lines for images, while higher-level areas handle more abstract information like object identity^10,11^. This mirrors our learning-phase results, in which early and higher-order sensory cortices were strongly engaged during perception. By contrast, retrieval runs the process in reverse, calling up and reactivating the stored representations instead^31^. Previous studies have shown that regions engaged during encoding are often reactivated during recall, particularly in associative sensory areas^4,12,13^. Our results support this, as we observed reactivation in associative auditory and visual cortices rather than in early sensory regions.

Additionally, we found deactivation in the early visual cortex during visual memory retrieval. Previous studies found that retinotopic populations in visual mnemonic and perceptual areas exhibit opposite responses during perception and memory recall^32–34^. This may explain the opposite responses in the early and associative visual cortex.

Both auditory and visual episodic memories were reconstructed in sensory association areas rather than early sensory regions. Previous fMRI studies have shown that the spatial pattern of neural activation for a specific visual episodic item during retrieval closely mirrors that during encoding, a phenomenon indicative of memory reinstatement^4–8^. In the present study, we replicated these findings in the visual domain using multivoxel pattern classification and ERS methods and, more importantly, demonstrated that auditory episodic memory items are reinstated in the higher-level auditory cortex. While several fMRI studies have reported that early visual areas can retain visual information during recall, these investigations typically used simple visual materials with only basic features, such as angles and lines, or deep neural network layers representing low-level visual features^35–40^. An EEG study by Linde-Domingo et al.^31^ found that higher-level conceptual information was first reconstructed before perceptual details were reconstructed during object memory recall, which is the opposite of the process during object perception. Thus, for complex dynamic visual and auditory scenes, the initial internal reconstruction involves higher-level item information represented in associative sensory areas, such as object category or scene meaning. This explains why episodic items were robustly reinstated only in these regions. Further studies are needed to confirm how different features of the same scene are hierarchically reinstated in sensory cortex.

Neural reinstatement measures from classification and PSI analysis offer an objective marker of memory quality that complements subjective vividness ratings. Prior studies have shown stronger visual memory reinstatement with higher vividness ratings^4–8^. Our study demonstrates that neural reinstatement could serve as a domain-general indicator of memory quality. With the same sample recalling auditory and visual episodes, visual and auditory memory vividness ratings are robustly correlated to sensory reinstatement. These results validate the reinstatement hypothesis by demonstrating that neural pattern reinstatement mediates successful memory retrieval across both auditory and visual modalities, thereby supporting its applicability to multimodal memory retrieval. This finding also indicates that sensory reinstatement measures could also serve as a potential marker for learning outcomes.

Our behavioural findings indicate that auditory is less vivid than visual episodic memory, even after additional training ensured that all participants memorized the stimulus pairs (Fig. 2A). This observation aligns with previous research showing that long-term auditory memory is generally less robust than long-term visual memory^1–3^. Gloede and Gregg^1^ found that when participants first memorize the pictures and environmental sounds, auditory representations are coarse and gist-based, while visual representations are highly detailed. Our findings support the idea that visual memory maintains a higher fidelity than auditory memory. However, balancing auditory and visual stimuli regarding perceptual salience and informativeness is difficult. Further studies must explore balancing the episodic sensory stimuli across modalities, enabling a more direct comparison of memory performance for balanced cross-modal stimuli.

The analyses of neuroimaging data reveal that, in addition to shared mechanisms, auditory episodic memory relies more heavily on internal representations than visual memory. Extending earlier behavioural work^1,2^, our multivoxel analyses suggest that visual memories retain finer perceptual details, whereas auditory memories depend more on abstract, gist-like representations. During visual memory retrieval, neural activity closely replicates the perceptual experience, reflecting high fidelity. Importantly, using internal-representation indices obtained from subtracting reinstatement and “reinstatement-free” residuals shows that auditory retrieval depends more on internally constructed representations, demonstrating that auditory memory is more abstract and gist-based. Therefore, listeners rely more on the internal construction of the sensory representation rather than faithfully encoding the auditory representation based on the external auditory stimuli. The informational interaction of memory reinstatement and representation between sensory association cortex and hippocampus subregions further supports these accounts. According to the hippocampal longitudinal gradient, the anterior hippocampal is more connected to the anterior temporal lobe and medial prefrontal cortex and represents the gist of an episode memory. In comparison, the posterior hippocampal is more connected to posterior sensory areas and represents the fine-grained perceptual details of an episodic memory^22,41^. We observed that the posterior hippocampus interacts exclusively with the sensory cortex during memory reinstatement, further supporting the idea that the reinstatement index reflects detailed sensory information about the recalled item.

In contrast, the anterior and posterior hippocampus interact with the sensory cortex during memory representation, with a greater coupling observed for the anterior hippocampus. Although this difference was not statistically significant, these preliminary results suggest that the representation index may be more closely associated with the gist-based aspects of recalled memories. In addition to the indirect evidence from RRS and hippocampal-cortical interaction, we performed categorical RSA to assess the gist-based representation directly. Gist representations of auditory items were better preserved from perception to memory than visual items. This result aligns with behavioural findings that auditory memory remains more enduring after several days^2^. It suggests that auditory memory may employ a strategy focused more on gist information than visual memory does.

The hippocampus plays a critical role in memory, with subregions contributing differently during retrieval^21,22^. While sensory reinstatement measures were significantly correlated with vividness ratings, we observed that the anterior hippocampus was specifically linked to these ratings. In contrast, the posterior hippocampus was uniquely associated with sensory reinstatement. This finding was unexpected if subjective vividness solely reflected the detailed information of memory, given that the posterior hippocampus is typically related to fine-grained details^22,41^. Previous research suggests that memory vividness does not always correspond to a high-fidelity replay of sensory details; instead, it may reflect the retrieval of an event’s general gist^42^. Thus, subjective vividness ratings and objective sensory reinstatement measures capture different facets of memory retrieval. Together, these results show that combining subjective and objective assessments provides a more comprehensive evaluation of memory quality.

Our results provide neural evidence that auditory and visual episodic memory retrieval partially show similar neural processing mechanisms. Both auditory and visual episodic memory are reinstated and robustly represented in the sensory association areas during memory retrieval. Auditory episodic memory is more gist-based and abstract, relying more on internal neural representation than the exact reactivation of encoding patterns. In contrast, visual episodic memory is more detailed, faithfully replicating the neural activation observed during encoding. These results provide insights that multimodal information is both redundant and, more importantly, complement to each other for us to better perceive and memorize the rich, complex sensory world around us.

## Methods

### Participants

The study included 25 participants (12 males, 13 females) aged 19 to 33 years old. The sample size is consistent with prior research^18,31,32^, and power analysis using G*Power indicates that it is sufficient to detect a medium effect size (0.5) with 79.8% power (paired t-test, one tail)^43^. Participants were recruited from the participant database of the Rotman Research Institute and provided written informed consent approved by the Rotman Research Institute’s Ethics Board. All participants had normal or corrected-to-normal vision, self-reported normal hearing, and reported no history of neurological or psychiatric disorders. Data from one participant was excluded because they did not complete the experiment.

### Stimuli

The stimuli comprised 16 natural or human sound clips and 16 natural or human video clips (for example, sound: human singing, wind blowing; video: human fighting, leopard chasing). Half of the clips depicted humanLJrelated content, and the other half depicted natural scenes with no humans. Each clip lasted six seconds. Two sequential stimuli formed a paired associate stimulus. The first stimulus (“cue”) served as a partial retrieval cue in a subsequent memory test. The second stimulus (“target”) was the target in the memory retrieval test. These paired associates included four auditory-auditory stimuli, four visual-visual stimuli, four auditory-visual stimuli, and four visual-auditory stimuli. In each case, the paired associate involved sequential presentation with no temporal gap between the leading and trailing stimulus. Pairings were pseudorandomized for each participant, while the cueLJtoLJtarget mapping was fixed: for 12 participants, Video 1–8 and Audio 1–8 served as cues and Video 9–16 and Audio 9–16 as targets; for the remaining 13 participants, this assignment was reversed.

### Procedure

Prior to the scanning sessions, participants engaged in the training phase to familiarize themselves with the experimental tasks (encoding and retrieval tasks). Four trials (one for each complex type) were presented using stimuli distinct from those used in the scanning sessions.

In the first scanning session, participants completed six encoding blocks and three retrieval blocks. One retrieval block followed two encoding blocks. In the encoding blocks, participants viewed/heard each of 16 paired associates in succession and were asked to memorize them. After each paired associate, participants rated the coherence of the cue and target from one (not coherent) to four (very coherent). In the test blocks, on each of the 16 paired associates, the retrieval cue (the 6s leading part) was presented, and participants mentally retrieved the target (i.e., the 6s trailing part). After retrieving, there was a 2-4 seconds (in 0.5-second increments) blank screen. Then, participants were asked to rate the vividness of their memory (1-4). Inter-trial intervals were 1-2 seconds (in 0.5-second increments).

After session 1, all participants received additional training until they could correctly identify every target when given its cue. In the second scanning session, each participant completed three encoding blocks and six retrieval blocks. Two retrieval blocks followed one encoding block. The learning and test blocks were the same as the first scanning session.

### Behavioural analysis

Four vividness ratings were averaged for each condition within each run. We performed repeated-measures analysis of variance to examine the effects of run order, cue modality, and target modality on vividness rating separately for the first scanning session and the second scanning session. A simple effect analysis was performed to determine if the interaction effect between the two factors was significant. One participant was excluded from the second session because no response was recorded under the audio-visual condition in the first block. Statistical analysis was conducted in R with the package bruceR^44^ and visualised using the package ggplot2^45^.

### Functional imaging data acquisition and preprocessing

Functional and structural MRI data were collected by a Siemens 3T Prisma MRI Scanner (Siemens Magnetom Trio) with a 64-channel head coil. T1-weighted images were acquired using the magnetization-prepared rapid acquisition gradient echo (MPRAGE) sequence (TR = 2000 ms, TE = 2.85 ms, field of view (FOV) = 256 × 240 mm, voxel size = 0.8 × 0.8 × 0.8 mm). Functional images were acquired using the multiband-accelerated echo planar imaging sequence (acceleration factor = 3, TR = 1390 ms for the first three participants, TR = 1400 ms for the other participants, TE = 25 ms, FOV = 220 × 220 mm, slices = 76, voxel sizes = 2 × 2 × 2 mm). Fieldmap images were acquired with the following parameters (TR = 4737 ms for the other participants, TE = 33.6 ms, FOV = 220 × 220 mm, slices = 76, voxel sizes = 2 × 2 × 2 mm).

All fMRI data were preprocessed using the standard pipeline of fMRIPrep 23.2.1^46^. Functional BOLD runs underwent slice timing, head-motion correction, susceptibility distortion correction (using fieldmaps), and co-registration to the T1-weighted reference. Time series were then normalized to the MNI152NLin2009cAsym template. Confound regression included framewise displacement, DVARS, global signals (CSF, white matter, and whole brain), and principal components derived from CompCor. Further details can be found in the fMRIPrep documentation.

For univariate analysis, additional preprocessing steps included spatial smoothing with a 6-mm full width at half maximum (FWHM) isotropic Gaussian kernel and scaling each voxel time series to have a mean of 100. The above steps were performed using the AFNI functions 3dmerge and 3dcalc^47^.

We used GLMsingle^48^ to estimate single-trial BOLD responses from unsmoothed, unscaled preprocessed fMRI data. All three processing stages generated robust single-trial beta estimates, including voxelwise HRF fitting using the library of HRFs, GLMdenoise, and fractional ridge regression. Following standard practice, the single-trial beta maps were extracted for multivoxel and trial-wise analyses.

### Univariate general linear model analysis

Single-subject multiple-regression modelling was performed using the AFNI program 3dDeconvolve^47^. Thirty-four regressors were defined, corresponding to cue and target presentations/recall for four conditions in both learning and test phases, each convolved with a 6s block function. Two regressors captured rating periods (coherence and vividness), each convolved with a 2s block function. In addition, motion parameters, CompCor components, global signal, and DVARS regressors were entered into the model to remove nuisance variance. Time points exceeding motion thresholds (0.5 mm) were censored.

Since this study focuses on sensory neural reinstatement of auditory and visual memory, we defined auditory and visual sensory ROIs based on HCP-MMP1 atlas^49^. All ROIs within ‘Auditory_Association’, ‘Early_Auditory’, ‘Temporo-Parieto-Occipital_Junction’,’Lateral_Temporal’, ‘MT+_Complex_and_Neighboring_Visual_Areas’, ‘Ventral_Stream_Visual’, ‘Early_Visual’, and ‘Early_Visual’ cortices were included in the univariate group analysis. Averaged beta values were extracted for each ROI. Paired t-tests assessed whether the activation was different under auditory memory retrieval and visual memory retrieval for each ROI. FDR correction was performed to adjust for multiple comparisons.

Note that all statistical tests used cue-corrected measures: for each auditory ROI, the auditory measure was defined as the difference between the auditory and visual values in that ROI, whereas for each visual ROI, the visual measure was defined as the difference between the visual and auditory values in that ROI.

Outliers, which are defined as elements more than three standard deviations from the mean, were replaced with the group mean. The same procedure was applied to the subsequent PSI, multivoxel pattern classification, and RSA analyses.

### ROI-based pattern specificity index analysis

We performed PSI analysis in ROIs that showed significant differences between audio and video memory retrieval. ROIs within early auditory cortex were also included (Figure 3A, Table S3). We calculated the correlation of the spatial activation pattern between the two trials for each condition. Then, we subtracted the between-item correlation coefficients from the within-item correlation coefficients. One-sided one-sample *t*-tests were used to examine whether the cue-corrected measures were greater than 0.

We examined whether neural patterns during memory retrieval reflected reinstatement of neural activity patterns during target encoding by correlating trial-wise retrieval data to the template pattern of the item during encoding, which is termed ERS. The template pattern of each item was the averaged spatial activity during the encoding phase. Then, we subtracted the between-item correlation coefficients from the within-item correlation coefficients to get the trial-wise PSI for reinstatement. The averaged PSI for each condition was calculated separately for the first and second scanning sessions within each sensory cluster.

PSI for EES was calculated by correlating trial-wise encoding data to the template pattern of the item during encoding. Here, the template pattern was calculated, excluding the run where the current trial existed. For example, when we correlated the trial-wise data of the first run to the template, the first run was excluded from the template calculation. PSI for RRS was calculated using the same process as target encoding, using retrieval data in the second session, which was consistent with ERS.

To investigate whether auditory and visual memory rely more on reinstatement or internal representations, we compared standardized measures derived from the auditory association cortex (auditory memory reinstatement and representation) and the visual association cortex (visual memory reinstatement and representation). Because differences, such as voxel count, between these regions can result in different measurement scales, the measures from each area were first standardized. We then computed an internal-representation bias score for internal representation by subtracting the reinstatement measure from the internal representation measure for each participant, separately for auditory and visual memory. A one-sample t-test was used to determine whether these bias scores differed significantly from zero. Finally, a paired t-test was conducted to compare the auditory and visual memory bias scores.

To further examine how reinstatement influences RRS, we calculated PSI in a “reinstatementLJfree” RRS analysis. First, for each ROI, we regressed the encodingLJphase template pattern of the corresponding item out of every retrieval trial. We then computed PSI for the residual (reinstatementLJfree) retrieval vectors using the same procedure applied in the original RRS analysis. Comparing the resulting reduction in RRS between auditory and visual memory using a two-sided paired t-test allowed us to quantify how strongly each modality depends on reinstatement.

### Trial-wise sensory reinstatement/representation-vividness rating correlation

A cumulative link mixed model was employed to examine the trial-wise relationship between sensory reinstatement measures and subjective vividness rating, since vividness rating is ordinal. Stimuli and cue modality were added as control variables. Random intercept was set for participants.

The same procedure was performed for the trial-wise correlation between sensory reinstatement measures and vividness rating. Linear mixed modelling with the same formula is also performed for illustration purposes. The above analysis was performed in R with the “ordinal” and “lme4” packages^50,51^.

### ROI-based representational similarity analysis

We performed RSA to examine categoryLJlevel (human vs. nonLJhuman) representations during memory retrieval. Bilateral temporal voice areas and fusiform face areas, which are crucial for processing human voices and faces, respectively^52,53^, were included in the analysis. Bilateral temporal voice areas were defined as the left and right temporal regions centred on peak coordinates identified by a Neurosynth metaLJanalysis using the term “voice.”

Representational dissimilarity matrices (RDMs; 1 − Pearson’s r) were computed from trialLJbyLJtrial correlations of activation patterns within these regions. The diagonal elements were set to zero and excluded from further analyses. Trials recalling the same item were averaged, yielding one 8 × 8 RDM for visual targets and another for auditory targets. Conceptual RDMs were constructed according to the human and nonLJhuman category assignment (see Fig. 6B). Each neural RDM was then correlated with its corresponding conceptual RDM (Pearson’s r) to assess how strongly category information was expressed in the voice and face areas. One-sided one-sample *t*-tests were used to examine whether the cue-corrected measures were greater than 0. Two-sided paired *t*-tests were used to examine whether the difference between the temporal voice area and the fusiform face areas, as well as the difference between encoding and retrieval within the temporal voice area and within the fusiform face areas, were greater than 0.

### Voxel-wise hippocampus-vividness rating/sensory reinstatement correlation analysis

First, we smoothed the trial-wise beta map with a 6-mm FWHM isotropic Gaussian kernel. Voxel-wise activation for each trial in the hippocampus was extracted for brain-behaviour correlation. For each participant, we correlated the trial-wise vividness rating and activation of each voxel in the hippocampus after controlling for stimuli and condition. Correlation analyses were performed separately for visual target retrieval and auditory target retrieval. The same procedure was also performed for correlating hippocampus voxel-wise activity with sensory reinstatement measures, i.e., averaged PSI and classification probability in auditory and visual association areas. Average correlation coefficients in the anterior and posterior hippocampus were extracted using Melbourne Subcortex Atlas^29^.

### Hippocampus-cortex informational interaction

MVPA and PSAS were also performed in the anterior and posterior hippocampus using the same procedure as described above to acquire trial-wise neural reinstatement and representation measures in the hippocampus subfield. Linear mixed-effect modelling was performed to examine the trial-wise informational correlation between the hippocampus subfield and sensory association areas. The variables are standardized for acquiring standardized beta coefficients. The formula is similar to the above equation except that the vividness rating was replaced with reinstatement or representation in the sensory association cortex. The anterior and posterior hippocampus were entered in the same model so that we could compare the standardized beta coefficients. We further use the Cumming’s procedure^30^ to test whether the standardized beta coefficients of the anterior and posterior hippocampus are statistically significantly different from each other. Firstly, we employed model-based parametric bootstrap for mixed models (bootMer function in LME4) with 2000 simulations to estimate 95% confidence intervals (CI) of the beta coefficient. Then, if the intervals overlap less than half the length of one CI arm, the two beta weights could be considered statistically significantly different from each other (p < 0.05)^30^.

## Acknowledgments

We thank B. Lau for helping with data collection.

## Data and materials availability

All data needed to evaluate the conclusions in the paper are present in the paper and/or the Supplementary Materials. The behavioral and fMRI data that support the findings of this study will be available in OSF.

## Supplementary Materials

### ROI-based multivoxel pattern classification analysis

We performed MVPC in the same ROIs as PSI analysis. The classifiers were trained using a support vector machine algorithm with a linear kernel. The cost parameter C was set to 1. The input features were univariate single-trial beta maps that were estimated using GLMsingle^41^. Classification analysis was performed in Python using scikit-learn library^43^. A one-tailed paired t-test was also performed here with the same purpose as PSAS.

Similar to PSI analysis, we trained the classifiers using target data from encoding blocks separately for each condition. The classifiers were then used to identify the recalled target among the stimulus set using neural activity from retrieval trials (encoding-retrieval classification). Averaged classification accuracy for each condition was calculated separately for the first and second scanning sessions within each sensory cluster. Trial-wise classification probabilities of the “true” item were also extracted for subsequent trial-wise analysis.

To investigate whether the targets were represented during the encoding (encoding-encoding classification) and retrieval phases (retrieval-retrieval classification), we employed leave-one-run-out cross-validation to evaluate classification performance using encoding data for each condition. Classification accuracy was also averaged for the first and second scanning sessions within each sensory cluster for the representation during the encoding phase. Regarding representation during the retrieval phase, we only analysed the second scanning session, consistent with encoding-retrieval classification analysis.

We employed leave-one-run-out cross-validation to evaluate classification performance using encoding data for each condition to investigate whether the targets were represented during the encoding phase. Classification accuracy was averaged for the first and second scanning session within each sensory cluster for the representation during the encoding phase. Regarding internal representation during the retrieval phase, the same process was performed using retrieval data in the second session.

A procedure analogous to that used in PSI was performed to test the bias for internal representation of visual memory in the visual association cortex and auditory memory in the auditory cortex, and compare the bias scores between auditory and visual memory. Also, retrieval-retrieval MVPC analysis was performed using the residual (reinstatementLJfree) retrieval vectors. Comparing the resulting reduction in retrieval-retrieval classification accuracy between auditory and visual memory using a two-sided paired t-test allowed us to quantify how strongly each modality depends on reinstatement.

**Table S1.** Statistics of MVPA measures. One-sided one-sample *t*-tests were used to examine whether the cue-corrected measures were greater than 0.

| Pattern specificity index |  |  |  |  |
| --- | --- | --- | --- | --- |
|  | Auditory association areas | Early auditory areas | Visual association areas | Early visual areas |
| EES | 5.80 (<0.001) | 4.71 (<0.001) | 7.00 (<0.001) | 12.17 (<0.001) |
| ERS | 1.75 (0.046) | 0.85 (0.202) | 3.85 (<0.001) | 0.88 (0.194) |
| RRS | 3.35 (0.001) | 1.29 (0.106) | 2.07 (0.025) | 0.77 (0.226) |
| Classification accuracy |  |  |  |  |
| EES | 10.81 (<0.001) | 14.06 (<0.001) | 16.23 (<0.001) | 18.63 (<0.001) |
| ERS | 2.09 (0.024) | 0.89 (0.192) | 4.48 (<0.001) | 0.64 (0.264) |
| RRS | 3.47 (0.001) | 0.99 (0.167) | 2.23 (0.018) | 0.87 (0.197) |

**Table S2.**
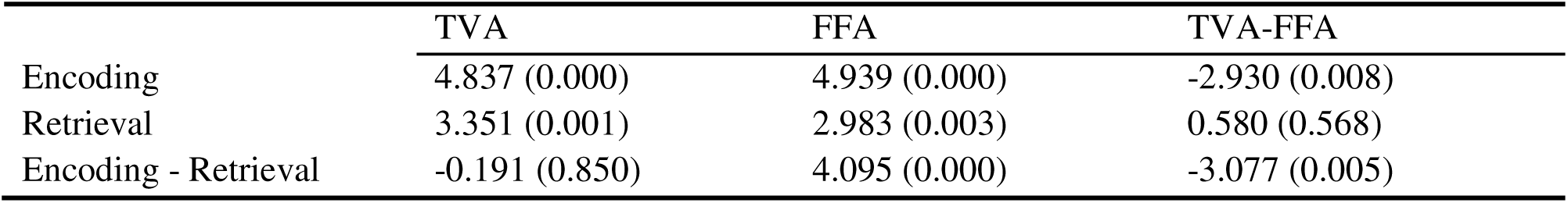
Statistics of RSA measures. One-sided one-sample *t*-tests were used to examine whether the cue-corrected measures were greater than 0 for encoding and retrieval in TVA and FFA. Two-sided paired *t*-tests were used to examine whether the difference between TVA and FFA and between encoding and retrieval within TVA and within FFA were greater than 0.

|  | TVA | FFA | TVA-FFA |
| --- | --- | --- | --- |
| Encoding | 4.837 (0.000) | 4.939 (0.000) | -2.930 (0.008) |
| Retrieval | 3.351 (0.001) | 2.983 (0.003) | 0.580 (0.568) |
| Encoding - Retrieval | -0.191 (0.850) | 4.095 (0.000) | -3.077 (0.005) |

**Table S3.** Hippocampal-cortical informational interaction analysis results. Standardized beta coefficient of anterior and posterior hippocampus predicting auditory and visual cortical memory reinstatement and representation from a linear mixed model.

|  | Visual association cortex |  | Auditory association cortex |  |
| --- | --- | --- | --- | --- |
|  | Visual representation | Visual reinstatement | Auditory representation | Auditory reinstatement |
| Anterior hippocampus | *B* = 0.099<br>P < 0.001<br>95%CI: 0.055, 0.144 | *B* = -0.005<br>P = 0.829<br>95%CI: -0.048, 0.039 | *B* = 0.055<br>P = 0.020<br>95%CI: 0.010, 0.100 | *B* = 0.040<br>P = 0.090<br>95%CI: -0.008, 0.085 |
| Posterior hippocampus | *B* = 0.046<br>P = 0.044<br>95%CI: 0.003, 0.091 | *B* = 0.065<br>P = 0.004<br>95%CI: 0.022, 0.109 | *B* = 0.044<br>P = 0.064<br>95%CI: -0.001, 0.088 | *B* = 0.074<br>P = 0.002<br>95%CI: 0.027, 0.120 |

**Figure S1.**
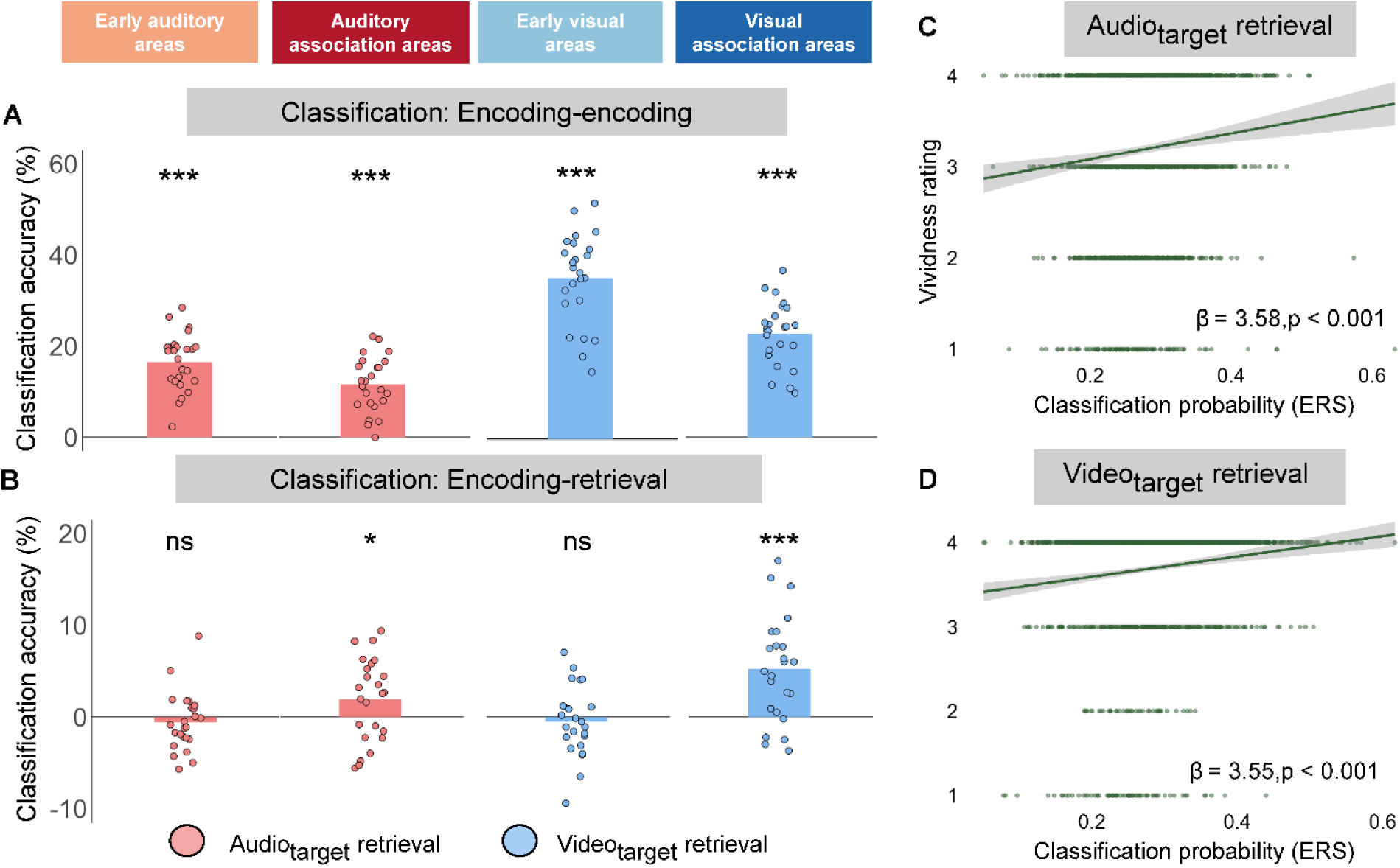
Neural reinstatement in sensory association cortex using MVPC analysis. (A) All four clusters showed robust representation of stimulus items during encoding. (B) Only visual and auditory association areas showed robust reinstatement of stimulus items during retrieval. (C, D) In both auditory and visual association areas, encoding-retrieval classification accuracy correlated with vividness ratings for their respective modalities. *P < 0.05, ***P < 0.001, ns, not significant.

**Figure S2.**
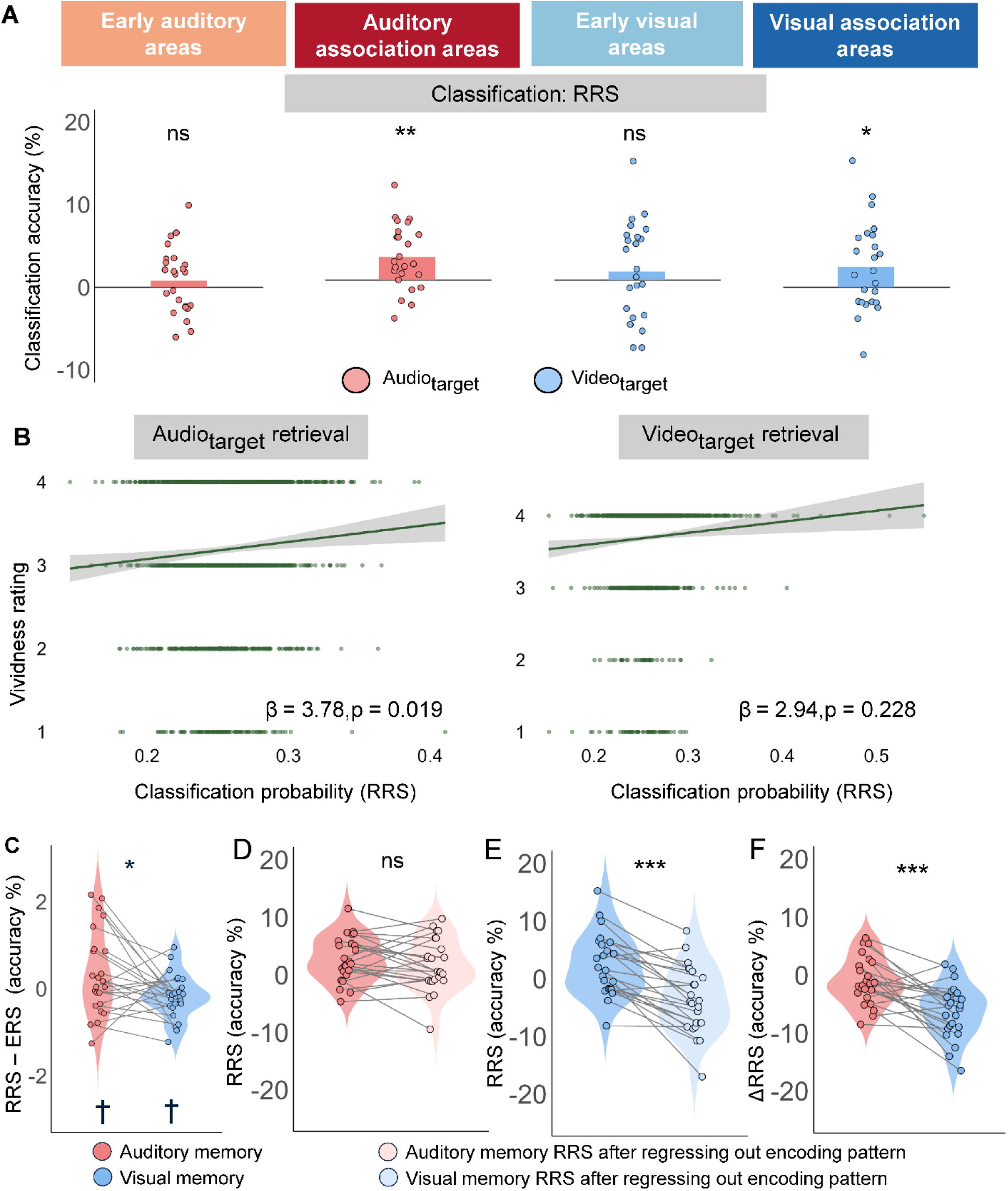
Internal representation in sensory association cortex using MVPC analysis. (A) Only visual and auditory association areas showed a robust internal representation of stimulus items during retrieval. (B) Auditory retrieval-retrieval classification accuracy correlates with vividness rating in auditory association areas. (C) Auditory memory showed a higher bias for internal representation than visual memory. (D, E) Regressing out trial-wise encoding patterns (to isolate activity unique to retrieval) reduced retrieval-retrieval classification accuracy only for visual modalities. (F) This reduction was smaller for auditory items in auditory association areas than for visual items in visual association areas. *P < 0.05, **P < 0.01, ***P < 0.001, ns, not significant.

**Table S3.** ROIs in neural reinstatement and representation analysis (Human Connectome Project multi-modal parcellation 1.0 (HCP-MMP1) atlas)

| Code | regionName | regionLongName | LR | Lobe | Cluster label in analysis |
| --- | --- | --- | --- | --- | --- |
| 23 | A1_L | Primary_Auditory_Cortex_L | L | Temp | Early_Auditory |
| 103 | RI_L | RetroInsular_Cortex_L | L | Par | Early_Auditory |
| 104 | PFcm_L | Area_PFcm_L | L | Par | Early_Auditory |
| 123 | PBelt_L | ParaBelt_Complex_L | L | Temp | Early_Auditory |
| 172 | MBelt_L | Medial_Belt_Complex_L | L | Temp | Early_Auditory |
| 173 | LBelt_L | Lateral_Belt_Complex_L | L | Temp | Early_Auditory |
| 203 | A1_R | Primary_Auditory_Cortex_R | R | Temp | Early_Auditory |
| 283 | RI_R | RetroInsular_Cortex_R | R | Par | Early_Auditory |
| 284 | PFcm_R | Area_PFcm_R | R | Par | Early_Auditory |
| 303 | PBelt_R | ParaBelt_Complex_R | R | Temp | Early_Auditory |
| 352 | MBelt_R | Medial_Belt_Complex_R | R | Temp | Early_Auditory |
| 353 | LBelt_R | Lateral_Belt_Complex_R | R | Temp | Early_Auditory |
| 102 | 52_L | Area_52_L | L | Temp | Auditory_Association |
| 106 | TA2_L | Area_TA2_L | L | Temp | Auditory_Association |
| 122 | STGa_L | Area_STGa_L | L | Temp | Auditory_Association |
| 124 | A5_L | Auditory_5_Complex_L | L | Temp | Auditory_Association |
| 128 | STSdp_L | Area_STSdp_posterior_L | L | Temp | Auditory_Association |
| 129 | STSvp_L | Area_STSvp_posterior_L | L | Temp | Auditory_Association |
| 138 | TPOJ1_L | Area_TemporoParietoOccipital_Junction_1_L | L | Temp | Auditory_Association |
| 174 | A4_L | Auditory_4_Complex_L | L | Temp | Auditory_Association |
| 302 | STGa_R | Area_STGa_R | R | Temp | Auditory_Association |
| 304 | A5_R | Auditory_5_Complex_R | R | Temp | Auditory_Association |
| 308 | STSdp_R | Area_STSdp_posterior_R | R | Temp | Auditory_Association |
| 309 | STSvp_R | Area_STSvp_posterior_R | R | Temp | Auditory_Association |
| 356 | TE1m_R | Area_TE1_Middle_R | R | Temp | Auditory_Association |
| 0 | V1_L | Primary_Visual_Cortex_L | L | Occ | Early_Visual |
| 3 | V2_L | Second_Visual_Area_L | L | Occ | Early_Visual |
| 4 | V3_L | Third_Visual_Area_L | L | Occ | Early_Visual |
| 5 | V4_L | Fourth_Visual_Area_L | L | Occ | Early_Visual |
| 6 | V8_L | Eighth_Visual_Area_L | L | Occ | Early_Visual |
| 21 | PIT_L | Posterior_InferoTemporal_complex_L | L | Occ | Early_Visual |
| 180 | V1_R | Primary_Visual_Cortex_R | R | Occ | Early_Visual |
| 183 | V2_R | Second_Visual_Area_R | R | Occ | Early_Visual |
| 184 | V3_R | Third_Visual_Area_R | R | Occ | Early_Visual |
| 185 | V4_R | Fourth_Visual_Area_R | R | Occ | Early_Visual |
| 186 | V8_R | Eighth_Visual_Area_R | R | Occ | Early_Visual |
| 200 | LO2_R | Area_Lateral_Occipital_2_R | R | Occ | Early_Visual |
| 201 | PIT_R | Posterior_InferoTemporal_complex_R | R | Occ | Early_Visual |
| 333 | VMV3_R | Ventromedial_Visual_Area_3_R | R | Occ | Early_Visual |
| 339 | VMV2_R | Ventromedial_Visual_Area_2_R | R | Occ | Early_Visual |
| 1 | MST_L | Medial_Superior_Temporal_Area_L | L | Occ | Visual_Association |
| 17 | FFC_L | Fusiform_Face_Complex_L | L | Temp | Visual_Association |
| 22 | MT_L | Middle_Temporal_Area_L | L | Occ | Visual_Association |
| 135 | TE2p_L | Area_TE2_posterior_L | L | Temp | Visual_Association |
| 136 | PHT_L | Area_PHT_L | L | Temp | Visual_Association |
| 137 | PH_L | Area_PH_L | L | Temp | Visual_Association |
| 139 | TPOJ2_L | Area_TemporoParietoOccipital_Junction_2_L | L | Par | Visual_Association |
| 140 | TPOJ3_L | Area_TemporoParietoOccipital_Junction_3_L | L | Occ | Visual_Association |
| 156 | FST_L | Area_FST_L | L | Occ | Visual_Association |
| 158 | LO3_L | Area_Lateral_Occipital_3_L | L | Occ | Visual_Association |
| 162 | VVC_L | Ventral_Visual_Complex_L | L | Temp | Visual_Association |
| 181 | MST_R | Medial_Superior_Temporal_Area_R | R | Occ | Visual_Association |
| 197 | FFC_R | Fusiform_Face_Complex_R | R | Temp | Visual_Association |
| 202 | MT_R | Middle_Temporal_Area_R | R | Occ | Visual_Association |
| 315 | TE2p_R | Area_TE2_posterior_R | R | Temp | Visual_Association |
| 316 | PHT_R | Area_PHT_R | R | Temp | Visual_Association |
| 317 | PH_R | Area_PH_R | R | Temp | Visual_Association |
| 319 | TPOJ2_R | Area_TemporoParietoOccipital_Junction_2_R | R | Par | Visual_Association |
| 320 | TPOJ3_R | Area_TemporoParietoOccipital_Junction_3_R | R | Par | Visual_Association |
| 336 | FST_R | Area_FST_R | R | Occ | Visual_Association |
| 338 | LO3_R | Area_Lateral_Occipital_3_R | R | Par | Visual_Association |

